# Metacognitive Introspection Alters the Dynamics of Pre-Decisional Neural Evidence Accumulation

**DOI:** 10.1101/2024.11.26.625501

**Authors:** Wei Dou, Shirin Afrakhteh, Jason Samaha

## Abstract

Metacognitive introspection refers to the capacity to monitor and evaluate one’s own performance, such as when expressing confidence in one’s choice. In the context of perceptual decision making, current computational models of confidence typically assume that confidence is computed after the evidence collection process underlying the initial choice has unfolded. This account predicts that the act of introspection required in order to provide a confidence judgment about one’s decision should have little impact on the dynamics of evidence accumulation leading up to that decision. We tested this by examining whether a neural proxy for perceptual evidence accumulation (the central-parietal positivity; CPP) exhibits different dynamics when participants do or do not rate confidence in their perceptual decisions. Behavioral results showed that confidence introspection increased decision accuracy and response time. Importantly, confidence introspection also resulted in a steeper CPP slope beginning shortly after (∼250 ms) decision evidence was available. We further observed an effector-specific neural signal of confidence evidence which both built up prior to the perceptual choice and which was predictive of the amount of confidence subsequently expressed by the participants. Our results indicate that introspection can alter the dynamics of perceptual evidence accumulation, possibly by acting on the gain of evidence accumulation, and that motor-related confidence signals emerge in tandem with pre-decisional evidence accumulation signals rather than post-choice.

## Introduction

When humans make perceptual decisions about the external world, they can typically introspect about the accuracy or uncertainty of their choices (Kepecs et al., 2008; Ratcliff, 1978). This metacognitive ability allows a decision-maker to predict and monitor the effectiveness of their performance and judge the probability that a decision will be correct. Metacognitive introspection is therefore thought to play a crucial role in guiding behavior, especially when external feedback is lacking (Shea et al., 2014). The underlying mechanics, as well as the functions of metacognitive introspection, are often studied by asking participants to judge their confidence in simple decisions (Deroy et al., 2016; Fleming & Dolan, 2012; Pereira et al., 2020). For instance, prior work has shown that people use their sense of confidence to determine whether adjustments to their decision-making strategies are needed in future events (Desender et al., 2018; Rollwage et al., 2020; van den Berg et al., 2016) and to rapidly correct decision errors in the absence of feedback (Yeung & Summerfield, 2012).

Increased attention to confidence in perceptual decision-making has led to the development of models aimed at understanding the fundamental computation involved in metacognitive judgments about perceptual decisions (Fetsch et al., 2014; Fleming & Daw, 2017; Kepecs et al., 2008; Kiani et al., 2014; Kiani & Shadlen, 2009; Maniscalco & Lau, 2016; Pleskac & Busemeyer, 2010; Sanders et al., 2016). Most current modeling frameworks make the often untested assumption that sensory evidence is first evaluated with respect to some decision boundary and, only after a decision is reached, confidence in that decision is computed (i.e., there is a sequential confidence assumption). For instance, in the framework of evidence accumulation, confidence is argued to based on the amount of evidence accumulated at the time of one’s choice and how long it took to accumulate that evidence (Dotan et al., 2018; Fetsch, Kiani, Newsome, et al., 2014; Geurts et al., 2022; Kiani et al., 2014; Kiani & Shadlen, 2009), implying that confidence is mapped from decision evidence leading up to the choice (i.e., pre-decision evidence) once the decision has been reached (see Fig. 1). Alternatively, post-decisional evidence accumulation accounts proposes that evidence continues to accumulate after the initial decision boundary is reached, and the accumulated post-decisional evidence is mapped onto a level of decision confidence (Desender et al., 2019, 2021; Resulaj et al., 2009; Van Zandt & Maldonado-Molina, 2004). Importantly, both pre-and post-decisional accounts seem to assume that confidence is the result of a mapping process that takes place only after the initial sensory evidence is accumulated and a decision is reached. Similar assumptions could be argued to exist implicitly in other frameworks, such as Signal Detection Theory or Bayesian accounts of confidence, which typically rely on evaluating confidence with respect to the initial decision boundary or the probability of that the initial choice was correct (Boundy-Singer et al., 2023; Geurts et al., 2022; Meyniel et al., 2015). However, these models typically do not have a temporal component so the strict ordering of computations is unclear.

**Figure 1.**
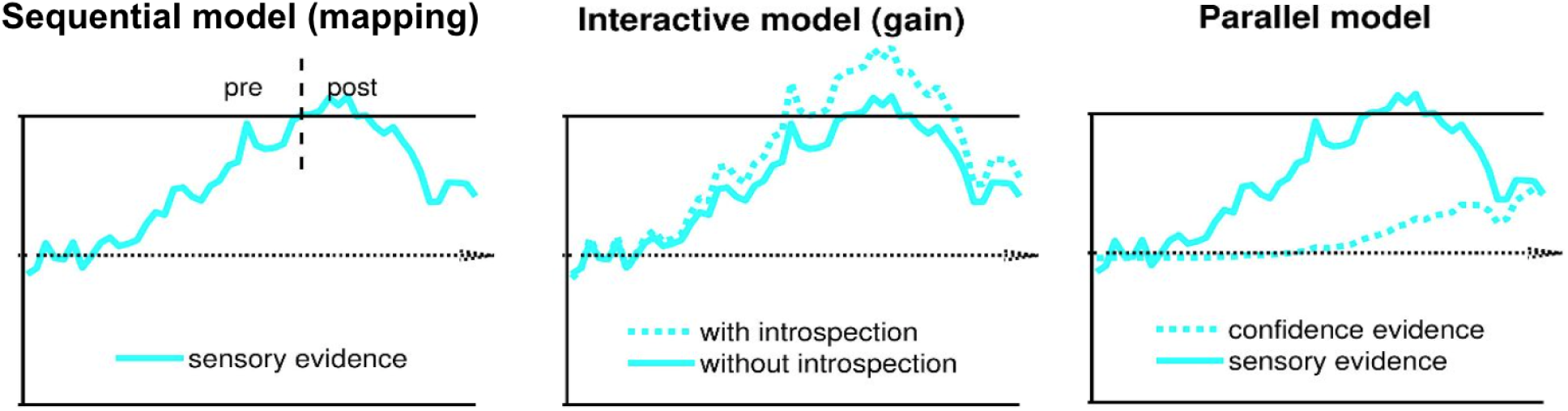
Possible evidence accumulation frameworks for how confidence relates to decision evidence. In each figure, the upper and lower horizontal solid lines represent the decision boundaries of two choices, and the middle horizontal dashed lines represent decision time. In the sequential choice-confidence framework (left), confidence is computed as a mapping from either pre-decisional or post-decisional sensory evidence (depending on the specific model) but only after sensory evidence is accumulated and a choice is reached. In the interactive model (middle), the act of introspection alters the process of sensory evidence accumulation before a first-order decision is even reached (shown here as a multiplicative “gain” change, although other forms of interaction are possible as well). As a result, sensory evidence is accumulated differently with introspection (dashed line) compared to without introspection (solid line). In the parallel model (right), sensory evidence for decision (solid blue trace) and confidence evidence (dashed blue trace) are accumulated in parallel. The parallel model is distinct from the sequential model in that confidence evidence is separately accumulated even prior to the first-order choice.

Here, we explored additional possibilities beyond the “sequential models” (as shown in Fig. 1, left) whereby sensory evidence is first accumulated and then mapped to confidence in sequence. Motivated by recent behavioral work suggesting that that computing confidence may not be sequential (Balsdon et al., 2020; Baranski & Petrusic, 2001; Double & Birney, 2024; Petrusic & Baranski, 2003), we formulated two additional accounts that could be tested at the neural level (Fig. 1). Balsdon et al. (2020) reported that an on-line sense confidence may control the evidence accumulation process for the first-order decision online by modulating the amount of sensory information necessary to commit to a decision. These findings suggest a different mechanism of confidence computation, which could be that confidence either alters, in some way, the trajectory with which sensory evidence is accumulated for decision-making (Interactive model; Fig. 1 middle) and/or that confidence accumulation unfolds in parallel with decision formation (Parallel model; Fig. 1 right), rather than after.

In the current study, we explored whether the act of introspecting (as when judging one’s confidence in a decision) would change the evidence accumulation process for decision making by combining recordings of electroencephalography (EEG) with decision behavior. The state of the accumulated evidence in our experiment was measured as the central-parietal positivity (CPP), an event-related potential (ERP) component thought to reflect sensory evidence accumulation given that the CPP build-up rate (or slope) correlates with evidence strength, reaction time, accuracy, and confidence (Dou et al., 2024; Kelly et al., 2021a; Kelly & O’Connell, 2013; O’Connell et al., 2012, 2018a). In our experiment, participants introspected their decisions in a random dot motion discrimination task on half of the trials by rating their confidence, but not on the other half of the trials. We discovered a steeper CPP slope for decisions accompanied by metacognitive introspection than for decisions without introspection, suggesting that evaluating confidence in one’s decision leads to enhanced sensory evidence accumulation, arguing against a strictly sequential model (see Fig. 1). Moreover, we found evidence for parallel confidence accumulation in a lateralized motor signal, specific to the effector used to report confidence, the build-up of which predicted participant’s confidence ratings prior to decision commitment.

## Methods

*Participants.* 32 participants (seven males, *M*_age_ = 19.65, *SD*_age_ = 1.34) were recruited from the University of California, Santa Cruz (UCSC) for course credit. They reported normal or corrected-to-normal vision and provided written informed consent. All procedures were approved by the institutional review board at UCSC.

*Stimulus and Apparatus.* A ∼53 cm wide electrically-shielded VIEWPixx/EEG monitor (120 Hz refresh rate, 1920 × 1080 pixels resolution) was used to present the stimuli on a black background. The distance between the monitor and a chinrest from which participants viewed the stimuli was ∼73 cm. Stimulus presentation and behavioral data were controlled by Psychtoolbox-3 (Kleiner et al., 2007; Pelli, 1997) running in the MATLAB environment.

The target stimulus comprised 150 white dots, each measuring 0.03°, displayed within a 5° circular aperture centered around a fixation point. For each stimulus, a certain portion of these dots, termed the “motion coherence level,” was randomly chosen for movement. These selected dots were displaced by a consistent 0.042° distance on each new frame, which corresponds to a motion speed of 5° per second either leftward or rightward (randomly selected on each trial). The remaining dots were placed randomly and independently within the circular aperture. Additionally, a red dot, measuring 0.25° in visual angle, was overlaid at the center of the dot stimuli, and participants were instructed to fixate on it. To discourage the tracking of individual dots and encourage responses based on the overall motion direction, each dot had a limited lifespan of 66 ms. Motion coherence was randomly selected on each trial from the following three levels with an equal likelihood: 4.5%, 20%, and 40%.

*Procedure.* Participants sat in a dim, sound-attenuated room and were asked to fixate on the central red dot on the screen throughout all trials. There were two main tasks involved in the experiment: the decision-only task and the decision-confidence task. Prior to starting each of the two tasks, participants first completed 180 trials of practice task using an easier set of coherence levels (5%, 30%, 40%, or 70%) than the main task. On each practice trial, participants were presented with a beep sound if they made an incorrect judgment about the motion direction. No feedback was provided during the main tasks. Participants advanced to the main task only if they attained an accuracy rate of 80% or higher on the 70% coherence practice trials. Participants who did not meet this standard in the initial practice block completed extra practice blocks until they achieved the required criteria. This criteria was established to ensure participants possessed a comprehensive grasp of the task and were capable of discerning the motion signals.

In the decision-only task, participants performed a random dot motion discrimination task (Kelly & O’Connell, 2013; Roitman & Shadlen, 2002) and were instructed to report the global direction of motion as quickly and accurately as possible (See Fig. 2A). At the start of each trial, a red dot appeared centrally for a random inter-trial interval between 1000 and 1500 ms. The dot motion stimulus was then presented for 300 ms, with the red dot remaining on the screen until participants hit a key to report the motion direction. Half of the participants were asked to place their left index finger and middle finger on the keys ‘S’ and ‘A,’ which were associated with rightward motion and leftward motion. The other half placed their right-hand index finger and middle finger on the keys ‘<’ and ‘>,’ which were associated with leftward and rightward motion.

**Figure 2.**
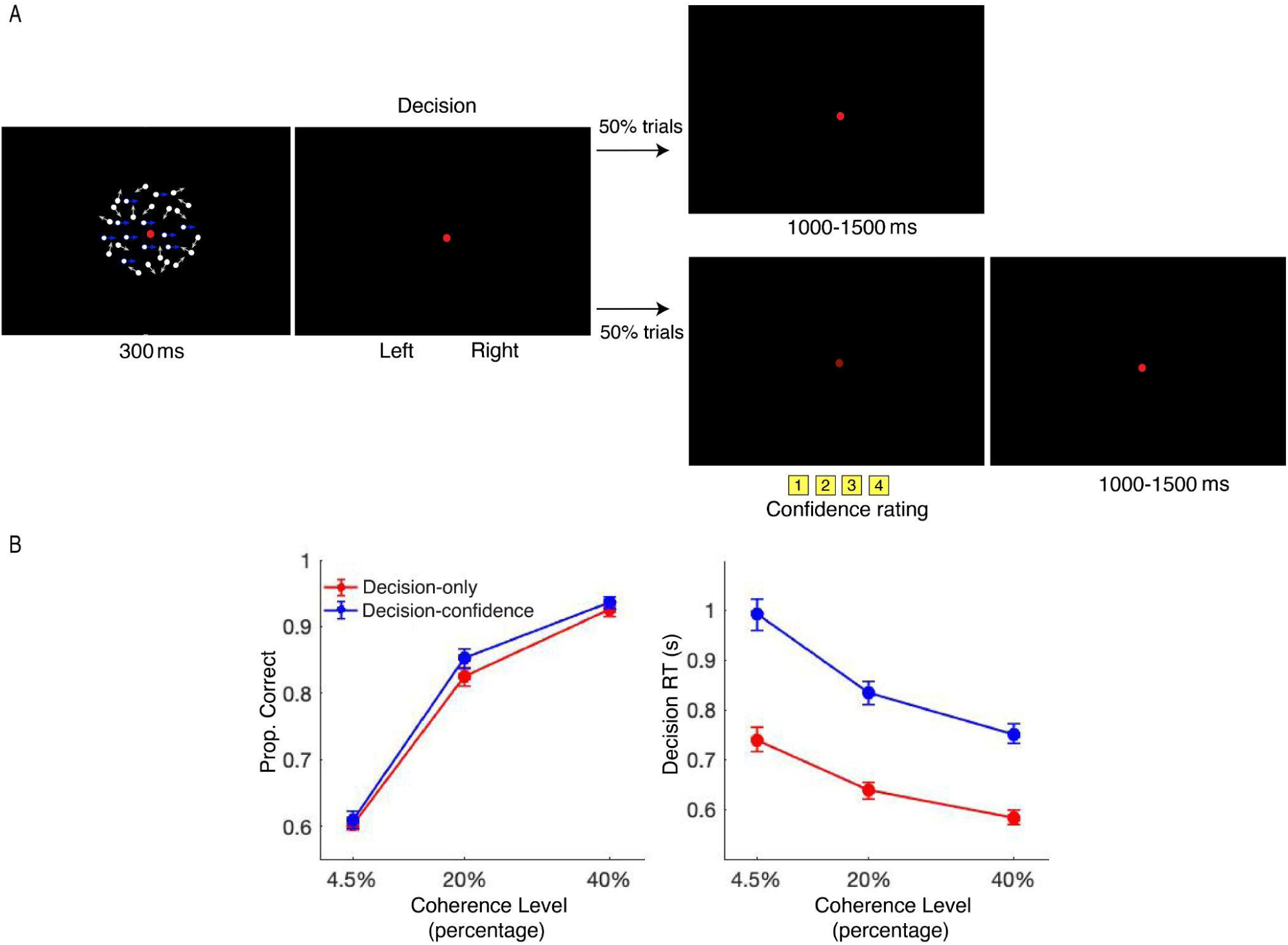
Example trial and behavioral results (N = 32). (A) Each trial started with a central red dot for 1000 to 1500 ms. A random dot motion (RDM) stimulus (motion coherence was randomly drawn from 4.5%, 20%, or 40%) then appeared for 300 ms. On 50% of the trials, participants performed the “decision-only” task, in which they only reported the motion direction (left vs. right) with one hand. On the other 50% of the trials, participants performed the “decision-confidence” task wherein participants first reported their decision (as in the decision-only task) but then also reported confidence in their decision on a 1-4 scale using their other (i.e., non-decision) hand. We hypothesized this manipulation would induce introspection about each decision. The two tasks were blocked, and their order of presentation was counterbalanced across participants. Half of the participants used their left hand for their motion direction choice and their right hand for confidence, and vice versa for the other half of the participants. (B) Motion direction discrimination accuracy (left) increased and RT of the motion decision (right) decreased with a higher coherence level. Moreover, in the decision-confidence task, accuracy was higher and RT was longer than in the decision-only task. Error bars denote ±1 SEM (across participants). Blue and white arrows in the RDM image represent example dot motion trajectories and were not shown in the actual displays.

In the decision-confidence task, after performing the same task described for the decision-only task, participants were asked to rate their confidence in their decision of motion direction on a 4-point scale. Participants were informed that “1” signifies a random guess at the motion direction and “4” signifies high confidence in their decision. The trial procedure was identical to the procedure used in the decision-only task, except that, once participants pressed a key to indicate the motion direction, the brightness of the central red dot decreased. The dimmer red dot remained on the screen until participants input a confidence rating. Participants used the same fingers as they used in the decision-only task to press the decision keys. Participants who pressed left-hand decision keys were directed to position their right-hand fingers (from index finger to little finger, respectively) on keys ‘J,’ ‘K,’ ‘L,’ and ‘;’ which corresponded to the confidence rating from 1 to 4. Participants who pressed right-hand decision keys were directed to position their left-hand fingers (from little finger to index finger, respectively) on keys ‘A,’ ‘S,’ ‘D,’ and ‘F,’ which corresponded to the confidence rating from 4 to 1.

Each participant completed 1080 trials in total, with 540 trials for each task, which consisted of 180 trials within each motion coherence level. The trials for each task were presented in three blocks, with 180 trials in each block. Motion direction and coherence level varied pseudo-randomly on a trial-by-trial basis. The two tasks were blocked, and the order of the presentation of the two tasks was fully counterbalanced across participants. Accordingly, 16 of the 32 participants performed the decision-only task first, with half of them using their right-hand fingers and the other half using left-hand fingers to press the decision keys. The remaining 16 participants performed the decision-confidence task first, with the position of the decision keys being counterbalanced between the right and left sides across the participants.

*EEG recording and analysis.* Continuous EEG was recorded by BrainVision actiCHamp with 63 active electrodes. The impedance at each central-parietal electrode was kept below 20kΩ. Recordings were digitized at 1000 Hz, and FCz was used as the online reference. EEG data was processed offline using custom scripts in MATLAB (version R2023b) and with EEGLAB toolbox (Delorme & Makeig, 2004). A high-pass filter at 0.1 Hz and a low-pass filter at 30 Hz were first applied to the EEG data using a zero-phase Hamming-windowed sinc FIR filter and then downsampled to 500 Hz. The data were re-referenced offline to the mean of all electrodes. Continuous signals were divided into epochs centered on stimulus onset, covering a time window from-2000 ms to 2000 ms. Trials were removed from EEG data if eye blinks were detected during stimulus presentation or if any scalp channel exceeded ±100 μV within the interval from-500 to 500 ms around stimulus onset. On average, 148 trials were removed per participant. These excluded trials were also not included in the behavioral analysis. Noisy channels were corrected using spherical interpolation, and an independent components analysis (infomax algorithm) was utilized to identify and remove one or two components per subject corresponding to eye blinks or eye movements. Finally, for each trial, we subtracted a pre-stimulus baseline ranging from-200 ms to 0 ms.

*Statistical analyses.* We computed the CPP build-up rate as the slope of a line fit to each participant’s average CPP waveform (Dou et al., 2023; Twomey et al., 2015) using a 200 ms sliding window advanced in 10 ms steps between 100 ms to 550 ms of the stimulus-locked CPP waveform and-1000 ms to-10 ms of the response-locked CPP waveform. Then separate linear models were fit to the data to predict CPP slopes at each time window by motion coherence level, RT (divided into 5 quantiles), accuracy (correct or incorrect), and confidence (high or low based on a participant-specific mean-split; decision-confidence task only). For each model, we compared the regression slopes at each time window to zero by a one-tailed t-test, hypothesizing that steeper slopes were associated with higher coherence, faster RT, correct trials, and higher confidence. To examine the effect of introspection on CPP, separate linear models were fit to the data in decision-only task and decision-confidence task separately. The regression slopes at each time window were compared using a two-tailed t-test, as we did not have a clear hypothesis about the task type effect. For all the analysis, a nonparametric, cluster-based permutation test was used to correct for multiple comparisons across time (Maris & Oostenveld, 2007). Specifically, we stored the largest cluster of significant time points across each of the 10,000 permutations of the data. Only clusters in the real data that surpassed the 95 percentile of this distribution of cluster sizes expected under the null hypothesis were considered significant.

## Results

### Preview

In the experiment, subjects viewed dot motion stimuli (300 ms) wherein a portion of dots moved coherently either to the left or right, while the remaining dots moved randomly. The portion of coherently moving dots (termed as motion coherence) varied between 2.5%, 20%, and 40%, which reflected the strength of motion signals. Participants (*N* = 32) performed two tasks across separate, counter-balanced blocks. In the decision-only task, participants made speeded judgements regarding the motion direction (left or right). In the decision-confidence task, in addition to reporting the decision about the motion direction, participants subsequently rated their confidence in their decision (Fig. 2). Our initial investigation focused on pinpointing a distinctive neural marker associated with accumulating perceptual evidence. This was accomplished by varying the evidence strength of the stimulus and evaluating the CPP in light of three characteristics observed in previous research (Kelly & O’Connell, 2013; O’Connell et al., 2012). Specifically, the CPP has been considered a neural signature of evidence accumulation, because its buildup rate shows (1) an increase with increasing evidence strength, (2) a decrease with long response times (RT), and (3) an increase in instances of correct choices relative to incorrect ones. We next explored whether there was a difference in the CPP slope between the decision-only and decision-confidence tasks.

### Judging confidence interferes with decision-making

A two-way within-subject ANOVA was applied to the behavioral data, with coherence (2.5%, 20%, or 40%) and task type (decision-only or decision-confidence) as the within-subject variable. As expected, increasing motion coherence led to higher decision accuracy (*F*(2, 62) =755.05, *p <.001*) and faster decision RT(*F*(2, 62) = 113, *p <*.001; Fig. 2B). Interestingly, there was a main effect of task type (decision-only versus decision-confidence) on accuracy (*F*(1, 31) = 4.48, *p =*.04) and RT (*F*(1, 31) = 86.69, *p <*.001), with the decision-confidence task leading to slightly higher accuracy (*M* =.85, *SE* =.01) and slower RT (*M* =.93, *SE* =.02) than the decision-only task (accuracy: *M* =.82, *SE* =.01; RT: *M* =.70, *SE* =.02). The interaction effect was significant on RT (*F*(2, 62) = 17.09, *p <* 0.01). Further analysis revealed that the RT difference between the decision-confidence task and decision-only task at 4.5% coherence level (*M* =.25, *SE* =.03) was larger than the difference at 20% coherence level (*M* =.20, *SE* =.02; *t*(31) = 3.29, *p* <.01), and the difference at 20% coherence level was larger than the difference at 40% coherence level (*M* =.17, *SE* =.02; *t*(31) = 3.38, *p* <.001). There was no significant interaction effect on accuracy (*F*(2, 62) = 1.88, *p =*.16). The results suggest that when task demands require introspecting about one’s decision, discrimination accuracy increases at the expense of longer RT. When considering the behavioral data alone, introspection could have possibly improved accuracy by either inducing a trade-off between speed and accuracy or by enhancing the process of evidence accumulation yet slowing down RTs due to additional pre-decision motor planning in service of the upcoming confidence report.

### A neural signature of perceptual evidence accumulation

During decision formation for both the decision-only task and the decision-confidence task, the EEG data contained a positive ERP component over central parietal electrodes which we refer to as the CPP component (O’Connell et al., 2012; Twomey et al., 2015) as its activity exhibited a discernible peak around 500 ms after stimulus onset (Fig. 3, *left* two columns) and an incremental build-up leading to the motor response (Fig. 3, *right* two columns). To further verify that this signal behaves as an evidence accumulation process for the motion decision, we examined the slope of the CPP component, which theoretically reflects the rate of evidence accumulation. A linear regression analysis was conducted on the average CPP waveforms of individual participants. The procedure entailed fitting a line to the waveform within a 200 ms sliding window, incremented in 10 ms steps. Separate linear models were then run to predict CPP slopes, at each time step, using motion coherence level, RT (divided into 4 quantiles), accuracy (correct or incorrect), and confidence (for the decision-confidence task only) as the predictors. To evaluate each model, we compared the regression slope at each time window against the null hypothesis of zero using a nonparametric cluster-based permutation test (Maris & Oostenveld, 2007).

**Figure 3.**
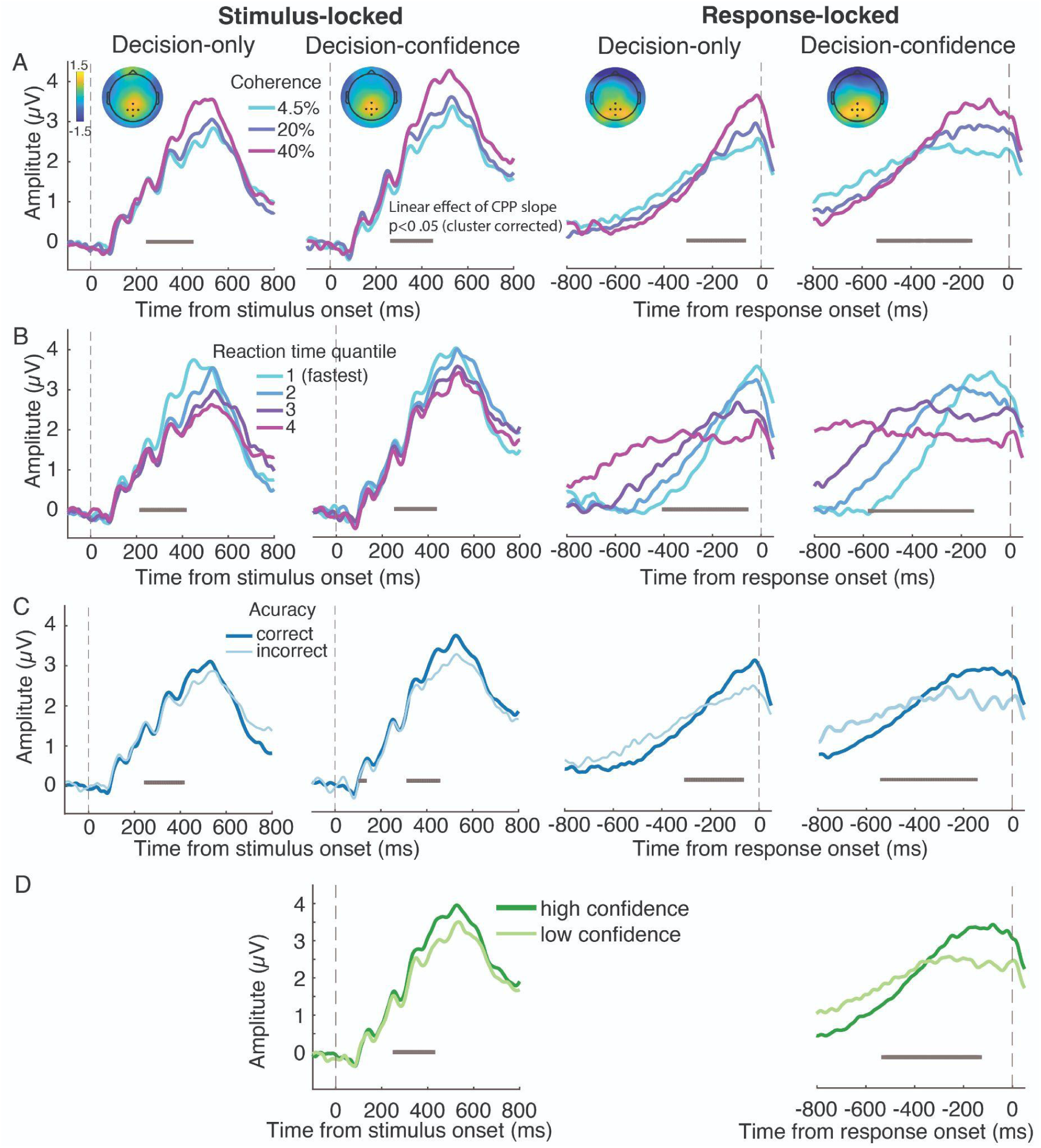
Grand average ERP time-courses locked to both the stimulus onset (left two columns) and response execution (right two columns). The vertical dashed lines denote stimulus onset (left two columns) and response execution (right two columns). (A) the inset topography of the ERP shows the difference in the spatial distribution of electrical activity between 40% and 4.5% coherence (40%-4.5%) after stimulus onset (stimulus-locked ERPs: 350 ms to 550 ms) and prior to response execution (response-locked ERPs:-200 ms to-10 ms). Black dots in the topographical plot highlight the central-parietal electrodes used in all subsequent analysis of CPP. Both stimulus-locked and response-locked CPP slopes increased as the motion coherence increased in both the decision-only task and the decision-confidence task. (B) In both tasks, stimulus-locked and response-locked CPP slopes also systematically steepened with faster RT (4 equally-sized RT bins). (C) In both tasks, both stimulus-locked and response-locked CPP slopes were higher on correct trials than on incorrect trials. (D) In the decision-confidence task, both stimulus-locked and response-locked CPP slopes were steeper when confidence was higher. Gray points below each ERP indicate the center of the 200 ms sliding time windows in which a linear effect of CPP slope as a function of coherence, RT, accuracy, or confidence reached significance ( p <.05) after correcting for multiple comparisons across time.

This analysis revealed that as participants judged motion signals with higher coherence (i.e., stronger evidence) both stimulus-locked and response-locked CPP slopes became steeper (Fig. 3A) in both tasks. Steeper CPP slopes were also observed when RTs were faster (Fig. 3B) and responses were correct (Fig. 3C) in both tasks, and higher confidence in the decision-confidence task (Fig. 3D). All the significant effects fell in the time range of 210 ms to 450 ms relative to post-stimulus and-590 ms to-50 ms relative to pre-response. The specific clusters in which a linear effect of CPP slope as a function of coherence, RT, accuracy, and confidence (p <.05) after correcting for multiple comparisons across time were denoted in each panel in Fig. 3. These results demonstrate that the slope of the CPP waveform in the data appears to encode accumulated evidence insofar as it predicts evidence strength, accuracy, RT, and confidence.

### The effect of introspection on the neural signal of evidence accumulation

Our main goal is to test whether introspecting one’s decision by judging confidence during the decision-making process could alter the way evidence is accumulated by comparing the CPP slope in the decision-only task with that in the decision-confidence task. As shown in Fig. 4, the CPP slopes in the two tasks were significantly different such that the decision-confidence task was associated with a steeper CPP spanning a significant cluster from 210 to 380 ms relative to stimulus onset (Fig. 4A left panel) and-800 to-540 ms relative to response onset (Fig. 4B left panel). The results also showed that the decision-confidence task was associated with a shallower response-locked CPP from-300 to-50 ms. Additionally, when the data were broken down by coherence levels, introspection had a significant effect on the slope of the stimulus-locked CPP at all levels of coherence (Fig. 4A right three panels) around 250 ms to 400 ms and the slope of the response-locked CPP at 40% coherence (Fig. 4A right panel) within-800 ms to-470 ms, consistent with introspection causing more rapid evidence accumulation. On the other hand, however, the slope of the response-locked CPP was shallower for the decision-confidence task at 4.5% and 20% coherence around-360 ms to-50 ms. Thus, while the stimulus-locked CPP shows a clear and consistent pattern of introspection causing steeper CPP slopes, the response-locked CPP shows a more complicated pattern with effects in both directions at different timepoints, a pattern we explain in the discussion. Interestingly, as can be seen in Fig. 4B, the peak amplitudes of the CPP prior to the motor response were similar in both tasks, which indicates a similar decision boundary may have been adopted for the decision in both tasks.

**Figure 4.**
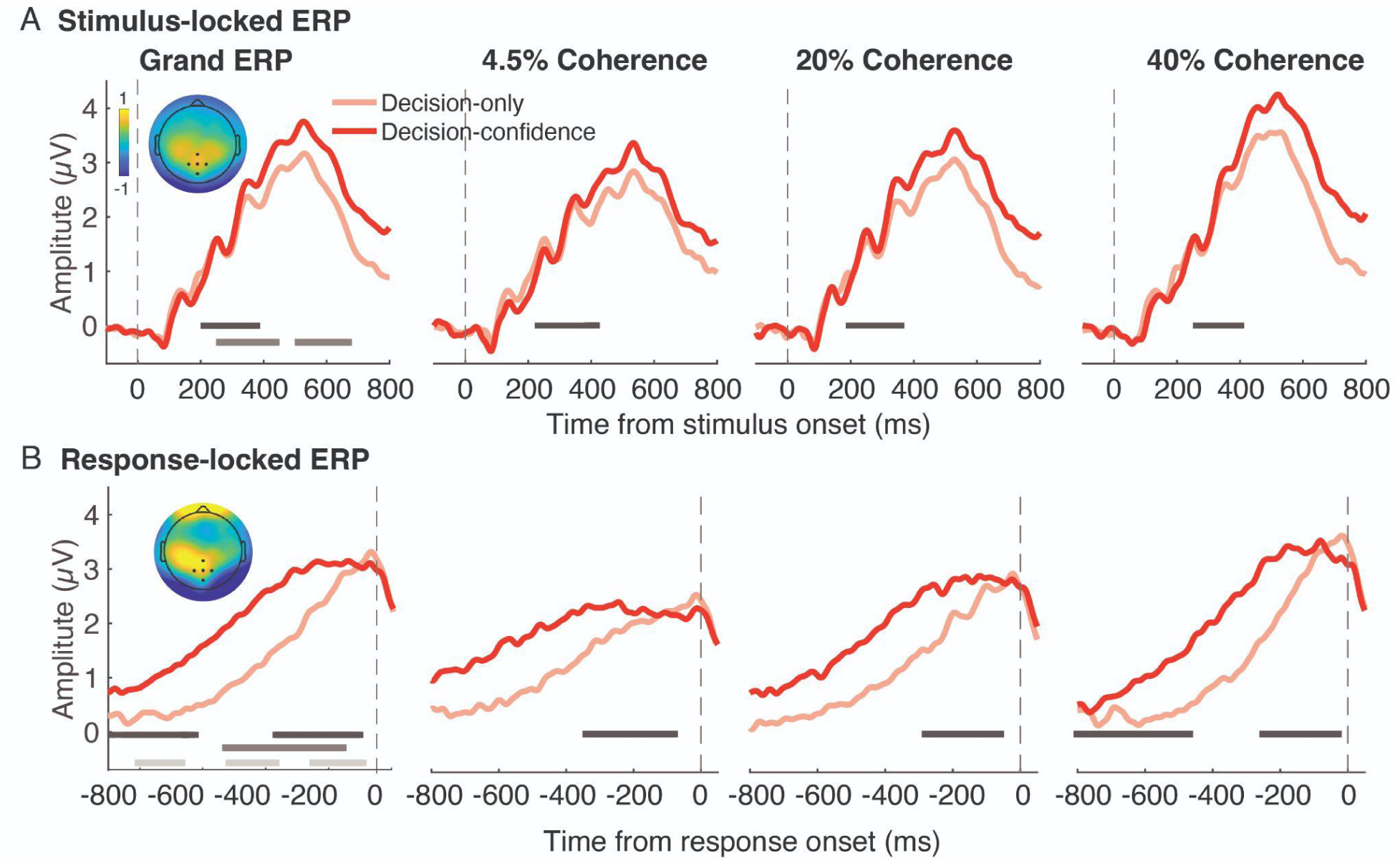
Grand average ERP time-courses locked to both the stimulus onset (A) and response execution (B). The inset topographies show the effect of introspection (decision-confidence minus decision-only) from 350 ms to 550 ms for the stimulus-lock and-250 ms to-50 ms for the response-locked ERP. (A) Stimulus-locked CPP slope was steeper in the decision-confidence task than in the decision-only task for the grand ERP (left panel) and for the ERPs at each coherence level (right three panels). (B) Response-locked CPP slope in the decision-confidence task differed from the slope in the decision-only task for the grand ERP (left panel) and the ERPs at each coherence level (right three panels). Black points below each ERP indicate the center of the 200 ms sliding time windows in which task type had a significant effect on the CPP slope (p <.05) after correcting for multiple comparisons across time. The dark gray points show a significant effect on coherence level, and the light gray points show a significant interaction between task type and coherence level.

When examining the topographical distribution of the effect of task type, particularly in the stimulus-locked analysis (see Fig. 4A), we noticed three distinct foci: one centered over the posterior midline where the CPP was maximal and also two additional possible effects over more anterior left and right motor areas. Given that the response hands were counterbalanced in our study, we wondered whether the left and right motor foci corresponded to the subset of participants who used their right or left hand, respectively, for the confidence response. This could provide evidence that an effector-specific confidence signal was emerging in parallel with the central parietal direction decision signal. To test this, we calculated the difference in both the stimulus-locked ERP and response-locked ERP between the two tasks (decision-confidence task minus decision-only task) for participants making right-hand confidence ratings (left-hand decisions) and left-hand confidence rating (right-hand decisions) separately (note that since participants are always making the direction decision with the same hand in both tasks, this subtraction cancels out motor activity related to the first-order decision). As shown in Fig. 5, the scalp topographies of those different ERPs exhibited clear lateralization within the time windows from 350 ms to 550 ms relative to stimulus onset and from-250 ms to-50 ms relative to response onset, such that the topography of the effect was lateralized over right central-parietal motor areas (electrodes: C4, C6, CP4, and CP6) for participants who made left-hand confidence ratings and over left motor areas (electrodes: C3, C5, CP3, and CP5) for participants who made right-hand confidence rating. Testing for task differences at these contralateral (to the confidence response) electrodes revealed that the slopes of the ERP components for both stimulus-locked and response-locked data were steeper in the decision-confidence task than in the decision-only task (see the time windows of the significant clusters denoted in Fig. 5). However, the ERP slope was steeper in the decision-only task from-210 ms to 0 ms relative to response onset in left-hand confidence rating participants (Fig. 5A). This may reflect faster evidence accumulation in the decision-confidence task, which reached the decision boundary earlier and began to decline due to leakage in the accumulation process, while evidence in the decision-only task continued to accumulate. This difference in the lateralized motor ERP, which was evident around 200 to 400 ms relative to stimulus onset and depended on which hand participants subsequently used for their confidence response, strongly suggests that confidence evidence, as reflected in confidence-effector-specific preparatory activity, begins before one’s perceptual decision is even reached.

**Figure 5.**
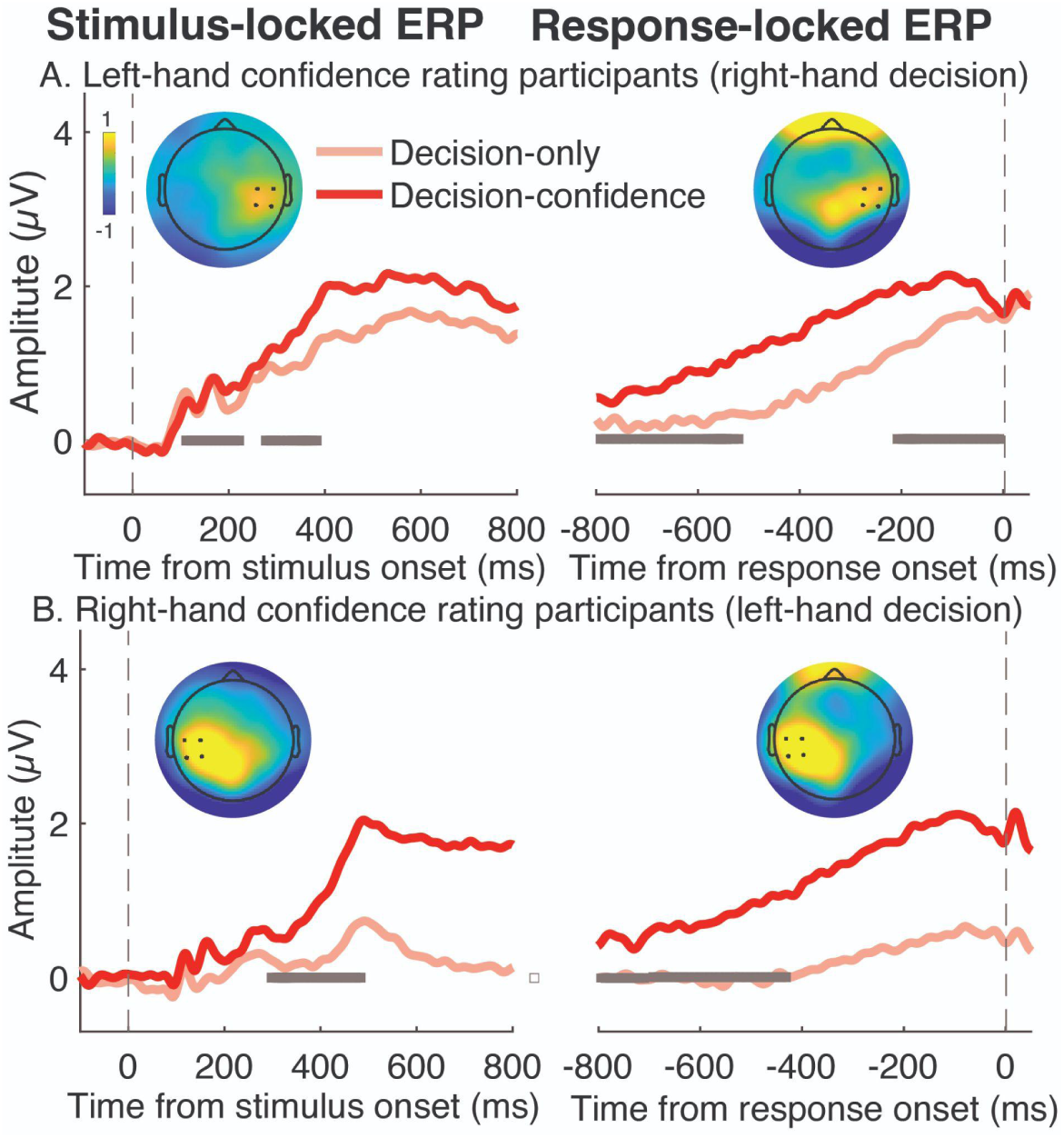
Grand average ERP time-courses of the decision-only task and the decision-confidence task locked to both the target onset (left column) and response execution (right column) for participants who made left-hand confidence ratings (A; N = 16) and right-hand confidence ratings (B; N = 16). The inset topographies show the introspection-related ERP difference (decision-confidence minus decision-only) from 350 ms to 550 ms for the stimulus-lock and-250 ms to-50 ms response-locked analyses. (A) The topographies show a positive-going right-lateralized component over motor areas contralateral to the confidence response (highlighted by the block dots which denote electrodes used in the analysis for participants who made left-hand confidence ratings). Both stimulus-locked and response-locked ERP slopes were steeper in the decision-confidence task compared to the decision-only task, except during the time window from-210 ms to 0 ms of the response-locked ERP, where the slope in the decision-confidence task was shallower. (B) The topographies show a positive left-lateralized component in participants who made right-handed confidence ratings. Both stimulus-locked and response-locked ERP slopes were higher in the decision-confidence task than in the decision-only task.

We next examined whether the pre-choice signal that emerged over sensors contralateral to the confidence hand could be used to predict participant’s confidence rating by contrasting contra-and ipsilateral motor signals for low and high confidence trials (using data from the decision-confidence blocks only). If the motor preparation signals represent the accumulation of confidence evidence, the slopes of ERP over contralateral motor areas should be associated with the confidence level. The results indicated that for participants who used their left hand to rate confidence (Fig. 6A), the slopes of the ERP recorded from contralateral electrodes significantly differed between high and low confidence within the time windows from 270 ms to 310 ms relative to stimulus onset and from-450 ms to-250 ms respective to response onset (Fig. 6A). For participants who used their right hand for confidence (Fig 6B), significant slope differences were observed within the time window from-240 ms to-170 ms relative to the response. In both groups, the slopes of ipsilateral stimulus-and response-locked ERPs also occasionally differed between high and low confidence trials, therefore we computed lateralized responses as well by subtracting the ipsilateral and contralateral responses to check if the steeper build-up on high confidence trials was indeed hemisphere specific. Indeed we found a significantly steeper lateralized build up rate on high confidence trials within the time window from 210 ms to 230 ms after stimulus onset and from-310 ms to-300 ms relative to response onset (Fig. 6C). Although the effect was brief, recall that these points reflect the center of a 200 ms window across which the waveform slopes are computed. These findings suggest that motor-related confidence evidence is being accumulated before a decision is made, in parallel with choice evidence.

**Figure 6.**
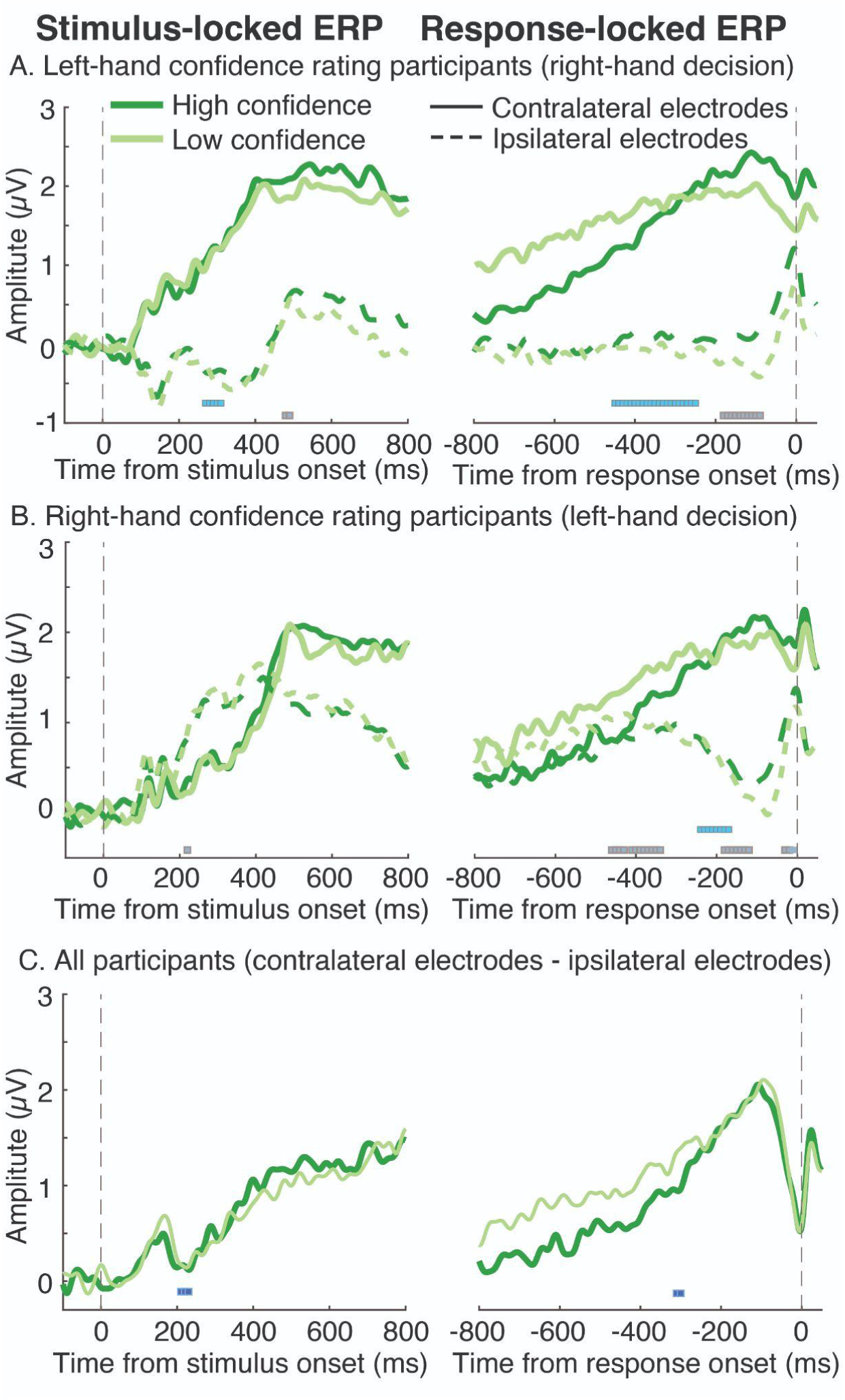
ERP time-courses in the decision-confidence task locked to both the target onset (left column) and response execution (right column). ERPs were extracted from the electrodes over motor areas contralateral (solid line) and ipsilateral (dashed line) to the confidence hand for participants (n=16) who made left-hand confidence ratings (A) and those (n=16) who made right-hand ratings (B). The slopes of response-locked ERPs were higher for high confidence than for low confidence. The light blue points show a significant effect of contralateral confidence-hand ERP on confidence, and the light gray points show a significant effect of ipsilateral confidence-hand ERP on confidence. (C) Lateralized ERPs were isolated by subtracting ipsilateral from contralateral waveforms (n=32). The dark blue points show a significant effect of confidence level on the lateralized ERP buildup rate.

## Discussion

The current study examined whether introspecting about one’s decision to provide a confidence judgment can alter the way that neural evidence is accumulated for that very decision. Behavioral results showed that confidence introspection led to higher decision accuracy and longer decision RTs, indicating a behavioral consequence of judging confidence. At the neural level, we first replicated previous findings showing that the CPP component indexes evidence accumulation as it is associated with evidence strength, decision accuracy, RT, and confidence judgments (Kelly et al., 2021; Kelly & O’Connell, 2013; O’Connell et al., 2012, 2018; Dou et al., 2023; Gherman & Philiastides, 2015; Herding et al., 2019; Vafaei Shooshtari et al., 2019). Critically, we found that the CPP slope was steeper when observers made decisions with, as compared to without, confidence judgments, an effect that emerged rapidly within a few hundred milliseconds from stimulus onset, suggesting a relatively early effect of introspection on decision dynamics, well before the initial decision was made. Interestingly, a motor response signal for confidence reports, which was sensitive to confidence level, was observed in parallel with evidence accumulation for the initial decision. These findings indicate that confidence is not merely a retrospective assessment of accumulated evidence for the decision but an interactive, parallel process that unfolds alongside sensory evidence accumulation.

A key implication of our findings is that confidence signals are computed “online” and can seemingly interact with sensory evidence accumulation processes. The consequences of real-time, interactive confidence computations for decision making remain to be fully understood, but our findings provide neural evidence to support recent behavioral findings suggesting that an online confidence signal can be used to control parameters of the decision-making process (Balsdon et al., 2020). Moreover, the fact that simply rating confidence in a choice can alter behavior and neural signals of evidence accumulation implies that studies which include confidence judgments in their decision tasks might be studying different underlying decision dynamics, leading to different model fits and behavior, for example, than those that do no include a confidence judgment. This could have implications for comparing the results of decision-making tasks across different experimental protocols.

More generally, our finding of a pre-choice effector-specific confidence signal challenges models of confidence that assume that confidence is computed sequentially, only after the initial decision process has reached a choice. This assumption is perhaps most pronounced in so-called *post-decisional* models of evidence accumulation, whereby only the evidence after the choice is made is considered for confidence (Boldt & Yeung, 2015; Desender et al., 2019, 2021; Resulaj et al., 2009). Even beyond post-decisional models, however, it is common to assume that a choice must first be made before confidence *in that choice* could even be considered. This manifests as models that define confidence as the state of the “losing” accumulator in a race between different choices (Kiani et al., 2014; Fetsch, Kiani, Newsome, et al., 2014), since which accumulator loses can only be defined after the “winning” accumulator reaches the decision boundary and a choice is made. Additionally, even models that propose that confidence is the mapping of pre-decisional evidence onto a probability of being correct propose that it is the state of accumulated evidence *once a decision is reached* (or when sensory evidence is terminated), that is used to compute the mapping to confidence (Zylberberg et al., 2016). Thus, our findings, coupled with recent work from our group showing that pre-decision EEG signals can be used to predict confidence even when controlling for evidence strength, RT, and accuracy (Dou et al., 2024), strongly suggest that confidence begins to emerge in tandem with evidence accumulation signals before a decision is made.

We interpret our results to clearly suggest that an effector-specific representation of confidence emerges in tandem with decision evidence, favoring a parallel account of confidence. We also suggest that this online representation of confidence can interact with evidence accumulation, as evident by the change in CPP slope during introspection blocks. However, one could argue against this latter point on the basis that the CPP signal could reflect a mixture of two separate processes: evidence accumulation and a secondary process related to metacognitive monitoring, rather than an interaction of monitoring on the evidence accumulation process. Although scalp signals cannot unambiguously rule out this possibility, we note that the topography of the introspection effect on the CPP (e.g., Fig. 4A) resembles that of the coherence effect on the CPP (e.g., Fig. 3A), which suggests that introspection is indeed interacting with evidence accumulation dynamics. This interpretation is also supported by the fact that there was an increase in accuracy and RT during introspection blocks (Fig. 2B), suggestive of some underlying change to the decision process itself. We speculate that introspection could be acting on the gain of sensory evidence accumulation, which would explain the steeper CPP slope and higher accuracy.

However, the finding of slower RTs with introspection is contrary to this interpretation, since enhanced evidence accumulation would be expected to hasten RTs. Examining the response-locked waveforms in Fig. 4B, however, reveals an interesting pattern in the confidence blocks whereby the CPP waveform plateaus prior to response execution and then remains elevated for several hundred milliseconds. This contrasts with the decision-only blocks where response execution occurs immediately after the CPP peak. The signal plateau associated with introspection could reflect an initial decision being made (the plateau point) and then the maintenance of information while preparing the confidence response, all before the initial decision is reported. This momentary lingering associated with introspection could be a natural explanation for the slower RT in the decision-confidence blocks and also explains why the effect of introspection on the CPP waveform goes in opposite directions at different time points (i.e., when the decision-confidence CPP waveform plateaus, the decision-only waveform is steeper). This CPP plateau could thus reflect a neural correlate of the non-decision time component of the total RT (here associated with preparing the confidence report; Ratcliff & McKoon, 2008; Ratcliff & Tuerlinckx, 2002). This view is in line with the work of Lei et al. (2020), who used functional magnetic resonance imaging (fMRI) to compare brain activity between perceptual decisions followed by confidence ratings and perceptual decisions followed by selecting a random digit. Their behavioral results mimicked those found in the currency study and the fMRI analysis revealed increased activation in the confidence rating condition in several brain regions associated with motor planning and metacognitive introspection, including the left supplementary motor area, left dorsal anterior cingulate cortex, left inferior frontal gyrus, and bilateral precuneus. The changes in brain activity in these regions were correlated with the changes in decision RTs between the two conditions.

In sum, our results reveal new dynamics underlying the relationship between decision-making and confidence at the neural level, suggesting that confidence signals emerge in tandem with and can influence the evidence accumulation process underlying perceptual decisions.

